# CRISPR/CAS9-MEDIATED LOSS OF VESICULAR GLUTAMATE TRANSPORTER IN SEROTONIN NEURONS OF THE DORSAL RAPHE NUCLEUS LEADS TO SYNAPTIC CHANGES AND ANXIETY-LIKE BEHAVIORS

**DOI:** 10.64898/2026.02.23.707446

**Authors:** Lydia Saïdi, Véronique Rioux, Marie-Josée Wallman, Sandeep Sundara Rajan, Emmanouil Metzakopian, Martin Lévesque, Christophe D. Proulx, Martin Parent

## Abstract

Vesicular glutamate transporter 3 (VGluT3) is expressed in a large subset of serotonin (5-HT) neurons of the dorsal raphe nucleus (DRN), suggesting a potential for glutamate co-transmission. Although VGluT3 has been implicated in the physiology of several non-glutamatergic neuronal populations, its specific role in the organization and function of 5-HT axons remains unclear. Here, we used CRISPR-Cas9 mediated knockdown and viral overexpression of VGluT3 in DRN 5-HT neurons of adult mice to assess its contribution to synaptic architecture in the lateral hypothalamus (LHA) and to 5-HT-related behaviors. VGluT3 depletion did not significantly alter synaptic incidence or organization of 5-HT DRN terminals in the LHA. In contrast, VGluT3 overexpression increased the proportion of asymmetric synapses without changing the overall synaptic incidence. In behavioral assays, VGluT3 depletion impaired motor coordination and increased anxiety-like, repetitive, and social behavior, whereas VGluT3 overexpression selectively reduced repetitive behavior. Basal locomotion and depressive-like behaviors were unchanged by either manipulation. Together, these findings indicate that VGluT3 modulates both the structural organization and behavioral output of DRN 5-HT neurons, supporting a modulatory role for VGluT3-dependent signaling within 5-HT circuits.

## Introduction

Glutamate-mediated neurotransmission requires the transport of glutamate into synaptic vesicles by members of the SLC17 family of transporters, which includes three vesicular glutamate transporters (VGluTs) ^1, 2^. VGluT1 and 2 exhibit complementary expression pattern in adult brain, VGluT1 being found in neurons of the cerebral cortex, hippocampus and cerebellum, and VGluT2 being abundant in many neurons of the thalamus and brainstem ^3–5^. Neurons whose canonical transmitter is glutamate typically express either VGluT1 or VGluT2 ^1, 2, 6, 7^. In contrast, the atypical VGluT3 is expressed by discrete neuronal populations that primarily release transmitters other than glutamate. VGluT3 has been reported in γ-amino-butyric acid (GABA)ergic basket cells of the cerebral cortex ^8, 9^ and hippocampus ^10, 11^, cholinergic (ACh) interneurons of the striatum ^9, 12–14^, serotonin (also known as 5-hydroxytryptamine, 5-HT) neurons of the dorsal (DRN) and median raphe nuclei (MRN) ^8, 9, 13–17^, peripheral sensory neurons ^18, 19^ and inner hair cells of the cochlea ^20,21^. Expression of VGluT3 in these populations suggests a potential for glutamate release along with their canonical transmitter. Indeed, co-loading of glutamate with ACh, 5-HT or GABA can modify vesicular quantal size ^22–25^ and leads to co-release of both neurotransmitters ^11, 26, 27^. This vesicular synergy is also observed in dopamine (DA) and noradrenergic neurons through VGluT2 ^28–30^, and in ACh, GABA and 5-HT neurons through VGluT3 ^11, 26, 27, 31–33^. The possibility that 5-HT neurons co-release glutamate was first suggested 30 years ago from patch-clamp recordings of postnatal rat 5-HT neurons in microculture ^34^. More recent studies confirmed fast glutamatergic transmission from striatal ACh interneurons and mesolimbic DA neurons ^35–38^. Although glutamate co-transmission has been described in several neuronal populations, the functional role of VGluT3 in 5-HT neurons of the DRN remains unclear.

The 5-HT system regulates behaviors including anxiety, mood, impulsivity, aggression, learning, reward and social interactions, and is implicated in affective disorders such as major depression, anxiety and obsessive-compulsive disorder ^39–43^. In mice, VGluT3 is expressed by most 5-HT neurons of the midbrain raphe nuclei ^16^, including the DRN and the MRN ^8, 9, 13^. The DRN houses the vast majority of the 26,000 5-HT neurons in the mouse brain ^44^. It can be divided into four major subregions along its rostro-caudal axis: dorsal (DRD), ventral (DRV), lateral (DRL), and interfascicular (DRI) parts ^45–47^. VGluT3 is not expressed uniformly across DRN 5-HT neurons. Most (≈ 80%) of 5-HT neurons in the DRV and DRI express VGluT3, compared with ≈ 25% in the DRD and only a small fraction in the DRL ^48^. At the single-cell level, our previous single-axon tracing study ^49^ along with the work of others ^47, 50–52^ highlighted the highly collateralized nature of DRN 5-HT axons, projecting extensively to widespread midbrain and forebrain regions. We also reported regional variations in VGluT3 content within 5-HT axons, suggesting complex trafficking mechanisms along these highly collateralized projections ^49^. Evidence for glutamate released from DRN 5-HT neurons has been reported in the hippocampus ^53^, orbital prefrontal cortex ^54^, nucleus accumbens (nAcc) ^55^, ventral tegmental area (VTA) ^56^, and basolateral amygdala ^57^. Nevertheless, not all 5-HT axon varicosities contain VGluT3 ^33, 49, 52, 56^. Therefore, at least two subtypes of 5-HT axon varicosities can be distinguished by VGluT3 presence or absence ^17^, and the proportion of these two subtypes varies across target sites ^49^. This heterogeneity suggests that 5-HT neurons can distribute vesicles with distinct transmitter contents across different collaterals. Such properties may contribute to the adaptive and functionally diverse nature of DRN 5-HT projections ^14^. Still, the physiological role of VGluT3 within 5-HT system remains to be clarified.

Genetic deletion studies in mice have revealed distinct roles for each VGluT. *Vglut1* and *Vglut2* constitutive knockouts are lethal, with VGluT1 deletion causing post-weaning death and VGluT2 deletion resulting in perinatal mortality ^58, 59^. In contrast, *Vglut3*-null (*Vglut3*-/-) mice are viable but show various neurological phenotypes. *Vglut3*-/- mice exhibit normal locomotor activity but seizure-like cortical discharges ^21^, congenital deafness due to impaired inhibitory circuit refinement ^60^, increased anxiety-like behavior ^27^, and altered nociception ^61^. These findings indicate that VGluT3 is involved in the function of several neuronal microcircuits, but they also highlight the limitations of constitutive knockouts for dissecting the role of glutamate released from specific neuronal populations, such as DRN 5-HT neurons. To address this, we used temporally and spatially controlled manipulations of VGluT3 expression in DRN 5-HT neurons in adult mice, using CRISPR/Cas9-mediated knockdown or viral overexpression. We then investigated how VGluT3 expression levels influence the morphology of 5-HT axon varicosities and assessed the contribution of VGluT3 expression to behavior. Our results indicate that altering VGluT3 expression modifies the morphology of 5-HT axon varicosities and that VGluT3 overexpression promotes the formation of asymmetric (putative excitatory) synapses in the lateral hypothalamic area (LHA). Behaviorally, VGluT3 depletion increases anxiety-like, repetitive and social behaviors.

## Materials and methods

### Mice

This study was carried out on 85 transgenic mice of both sex (ePet-Cre;Cas9-eGFP) aged of 8-10 weeks old and weighing between 20-30g. They were obtained by crossing ePet-cre mice (transgenic B6.Cg-Tg(Fev-cre)1Esd/J allele) that express the Cre recombinase under the *pet1* promoter ^62^ with Rosa26-LSL-Cas9 knockin mice, which have a floxed-STOP cassette preventing the expression of downstream *Cas9* and *eGFP* sequences. Both mouse lines were purchased from Jackson Labs (stock #012712 and #024857, respectively). Two WT CD1 mice of 10 weeks old were employed as target mice for the social interaction behavioral test.

Mice were housed in group under a 12h light-dark cycle with water and food ad libitum. All procedures were approved by our Institutional Animal Care and Use Committee (*Comité de Protection des Animaux de l’Université Laval,* VRR-2022-1186), in accordance with the Canadian Council on Animal Care’s Guide to the Care and Use of Experimental Animals. Maximum efforts were made to minimize the number of animals used. Individuals in charge of behavioral measurements and data analysis were blind to experimental conditions.

### *Slc17a8* dgRNA validation

Two gRNAs against *Slc17a8* (*Vglut3* gene) were designed. The first gRNA targeted exon 1 and the second one targeted exon 2. They were cloned into gRNA expression plasmid (pLVPB-U6gRNA(Bbs1)-PGKpuro2ABFP). This plasmid containing the 2 gRNAs was co-transfected along with a puromycin-resistant plasmid encoding Streptococcus pyogenes Cas9 (SpCas9) (Addgene, plasmid #62988) in mouse fibroblasts NIH-3T3 (ATCC, CRL-1658) using lipofectamine LTX (Invitrogen). Co-transfection of the puromycin-ScCas9 plasmid along with a Cas9 activity reporter plasmid (Addgene, plasmid #67982) was used as a control condition. Three days post-transfection, cells were passaged and treated for three days with 2.0 µg/ µL puromycin. The cells were recovered for another two weeks, and genomic DNA was extracted using Monarch® Spin gDNA Extraction Kit (New England Biolabs, T3010). The exons 1 and 2 of *Slc17a8* have been amplified by PCR using Q5® High-Fidelity DNA Polymerase (New England Biolabs, M0491) using primers identified in Table S1. The PCR amplicons were sequenced by SANGER at the sequencing platform from CHU of Laval University. The spectrum of nucleotide Insertions and Deletions (INDELS) and the frequency of aberrant base pairs following the expected cut site were measured using Tracking of INDELS by Decomposition (TIDE) software, https://tide.nki.nl/ ^63^.

### AAV viral particle preparation

pAAV U6-dgRNA(Scramble)-hSyn-DIO-mCherry, pAAV U6-dgRNA(VGluT3)-hSyn-DIO-mCherry, pAAV CAG-DIO-VGluT3-P2A-mCherry and AAV serotype 2/DJ viral particles were produced by the Canadian neurophotonics platform viral vector core (POM).

### Stereotaxic injections

Two-month-old ePet-cre;Cas9-eGFP mice were anesthetized with 2% isoflurane and their heads were secured in a stereotaxic apparatus. Using a nanoinjector, a 1μL volume of virus [AAV-dgScr (1.2E13 GC/ml), AAV-dgVGluT3 (1.5E13 GC/ml) or AAV-VGluT3 (8E12 GC/ml)] was injected into the DRN using a 32° angle from the coronal midline at the following coordinates: (AP: -4.45/-4.65mm relative to bregma; ML: 2.0 mm; DV: -2.75/-3.50 mm).

### Behavioral Assessment

Four weeks following AAV injections, mice were subjected to a panel of behavioral tests as described below to assess any motor deficit, anxiety, behavioral despair, and anhedonia-like states using the open field, rotarod, nestlet shredding test, elevated plus maze, tail suspension test, social interaction test and sucrose preference test.

#### Open field

The locomotor activity of AAV-injected mice was assessed in an open field arena (42 x 42 x 30 cm), framed by transparent plexiglas walls and equipped with 16 light beams located in both the horizontal and vertical planes (AccuScan Instruments, Inc.). The activity was monitored using the activity-monitoring VersaMax version 3.0 software (AccuScan Instruments, Inc.). Mice were habituated to the room for 1 hour and were tested over a 60-minute period in the open field. Activity was measured by recording stereotypical and vertical activity and was quantified as the number of stereotypical and vertical movements, respectively. Total distance travelled during 60 min was measured as a locomotor activity parameter. Time spent in the center of the arena was also measured and used as an indicator of anxiety-like behavior.

#### Rotarod

Motor coordination was tested in AAV-injected mice using a Roto-Rod System (IITC Life Sciences, California, USA), with the use of an accelerating rather than constant speed protocol that was shown to minimize interference from learned memory ^64^. A habituation trial was done 1 week before, at constant speed, for 5 minutes. Rotations started immediately after mice were placed on the rotarod and accelerated from 4 to 40 RPM over a period of 300 s. Latency to fall (s) was recorded on 3 trials to limit the effect of motor learning. Group means were obtained from the best performance of the 3 trials.

#### Nestlet shredding test

Mice were removed from their home cages and placed individually into a fresh standard mouse cage with clean bedding and a standard cotton nestlet square (5 cm × 5 cm, 3 g). The nestlets were pre-weighed before the start of testing to calculate the % shredded at the end of the task ^65^ and evaluate the mouse’s anxiety level. Mice were left undisturbed for 30 min, after which they were returned to their home cages. The weights of the remaining non-shredded nestlet material were recorded, as previously described ^65^.

#### Elevated-plus maze test

Mice were put in the middle of an elevated-plus maze (EPM) apparatus, placed on a pedestal located at 50 cm above floor level, under red light condition. Each arm of the maze measured 50 x 10 cm. The black Plexiglas cross-shaped maze consisted of two open arms with no walls and two closed arms (40 cm high walls). Anxious behavior was assessed using an automated system (ANYmaze, version 5.0) and measured as the total time spent into the open arms.

#### Tail suspension test

The tail suspension test was adapted from a published protocol ^66^. Mice were habituated to the experimental room for 1 hour then suspended by their tail for 6 min. Clear hollow cylinders (4 cm length, 2.3 cm outside diameter, 2.2 cm inside diameter, 1.5 grams) were placed around their tail to prevent tail climbing behavior. A camera (GoPro Hero v.5) was placed in front of the tail suspension box (35 height X 74 width X 12 cm depth) for video recording. During the behavioral analysis, the time that each mouse spent as mobile was measured. The immobility time was reported to show the escape-related mouse motivation level.

#### Social interaction test

Mice were transferred to a quiet room under red-light condition and were habituated for 1 hour. For the first session, the subject animal was placed in an open-field arena with a small, wired enclosure on one side of the arena. The exploration behavior was tracked from above using the video camera connected to a computer running ANYmaze tracking software (version 5.0). In the first session, the mouse was allowed 2.5 min to explore the empty arena. In the second session, a WT C57BL/6 target mouse (of the same gender) was placed in the small enclosure of the arena, and the subject mouse was placed back in the arena for another 2.5 min. Social interaction was assessed by calculating the SI ratio, which is the amount of time that the animal spent in the interaction zone while the target mouse was present, over the time spent in the interaction zone while the target mouse was absent. The subject mouse behavior was considered to show a depressive-like behavior if the SI ratio was lower than 1.

#### Sucrose preference test

The sucrose preference test used in the present study was adapted from a published protocol^67^. Briefly, mice were given two bottles filled with water for a 24-hour habituation period. The following day and during the next 24 h, one of the two bottles was filled with a 1% sucrose solution. The two bottles were then weighed, and the position was switched for an additional 24 h. The total duration of the test was 48 h. Sucrose preference was calculated by determining the sucrose solution consumption, divided by total liquid consumption.

### Tissue Preparation

Six weeks following AAV injection, mice were deeply anesthetized with ketamine-xylazine (10 mg/mL and 1 mg/mL, respectively) and transcardially perfused with ice-cold sodium phosphate-buffered saline (PBS, 0.1 M; pH 7.4) followed by 4% paraformaldehyde diluted in 100 mM phosphate buffer, pH 7.4. Brains were extracted and post-fixed in the same solution for 24h at 4 °C, and then cut with a vibratome (Leica, model VT1200S) into 50 μm-thick coronal sections, which were serially collected in PBS. Sections were kept in an antifreeze solution at -20°C until immunostaining.

### Immunohistochemistry

The mouse brain sections were incubated in a blocking solution composed of 2% normal serum and 0.1 % Triton X-100 diluted in PBS for 1 h at room temperature (RT) and subsequently incubated overnight at 4 °C with primary antibodies at optimized concentrations. The following primary antibodies were used: chicken anti-GFP (1:1000; GFP-1020; Aves Labs); rat anti-mCherry (1:500; M11217; Invitrogen); rabbit anti-VGluT3 (1:500; 135 203; Synaptic Systems); sheep anti-TpH (1:250; AB1541; Millipore). Appropriate secondary antibodies conjugated to Cy5 (711-175-152; Jackson ImmunoResearch), Alexa-Fluor 488 (703-545-155; Jackson ImmunoResearch), Alexa-Fluor 594 (A-11007; Invitrogen) or Alexa-Fluor 647 (A-21448; Invitrogen) were used (1:400). Immunstained sections were mounted and coverslipped with fluorescence mounting medium (DAKO).

### Fluorescence in situ hybridization

One section per mouse (4 mice per group) was taken at the mid rostrocaudal level of the DRN and used for fluorescence in situ hybridization. RNAscope probes for mouse VGluT3 (Slc17a8) and mCherry (mCherry-C2) were designed by Advanced Cell Diagnostics (Newark, CA, USA). In situ hybridization was performed using the RNAscope Multiplex Fluorescent v1 assay according to the manufacturer’s instructions. The sections were heated and treated with protease digestion followed by hybridization with a mixture containing specific target probes for mouse Slc17a8 and mCherry mRNAs. mRNA for Slc17a8 was labeled with Alexa Fluor-488 whereas mCherry was detected with Atto-550. Sections were mounted on microscope slides and coverslipped with DAKO.

### Confocal imaging

Images obtained from immunostained sections were acquired with a confocal microscope (LSM 700, Zeiss) using 10x, 20x or 63x objectives, with constant acquisition parameters across sessions and experimental conditions. Counting of immunostained cells located in the DRN was performed by using the ImageJ software on tile scans (20x, 3 x 3) obtained from 3 equally-spaced sections per mouse (4 mice per group) taken at -4.36, -4.60 and -4.84 mm relative to the bregma^68^. An average number of 838 ± 204 TpH+ cells, 645 ± 115 GFP+ cells and 536 ± 65 mCherry+ cells were counted in the whole DRN of each mouse. To assess the presence and amount of a given mRNA or protein in DRN neurons, the mean fluorescence intensity for that channel was obtained in each mCherry+ cell body by using the ImageJ software. In the lateral hypothalamic area (LHA), to estimate the number of 5-HT axon varicosities per 10 μm immunoreactive for VGluT3, about 200 mCherry+ varicosities were counted in each mouse (3 sections per mouse, 4 mice per group), and the length of mCherry+ axons examined was determined from 40 µm-thick z stacks (40x, 0.5µm intervals) along their orthogonal views (Fig. S2B) using the Fiji/Image J software (https://fiji.sc) and the Zeiss LSM Viewer software (http://www.zeiss.de ⁄ downloads; version 3.5).

### Electron microscopy

For electron microscopy, one section per mouse (4 mice per group) taken at -2.30 mm from the bregma according to the mouse brain atlas of Franklin and Paxinos ^68^ was immunostained for mCherry as described above, but in the absence of Triton X-100, which was replaced by 0.5% gelatin. Sections were osmicated, dehydrated in ethanol and propylene oxide, and flat-embedded in Durcupan ACM (catalog no. 44611-14; Fluka) to be processed. Quadrangular pieces taken from the LHA were cut from the flat-embedded mCherry-immunostained sections. After being glued on the tip of resin blocks, they were cut ultrathin (80 nm) with an ultramicrotome (model EM UC7, Leica). Ultrathin sections were collected on bare 150-mesh copper grids (G150-Cu; Electron Microscopy Sciences) and examined by using a transmission electron microscope (Tecnai 12; Philips Electronic), at 100 kV, and an integrated digital camera (MegaView II; Olympus). The mCherry+ varicosities were sampled randomly, at a working magnification of 11,500x, by taking a picture of every such profile encountered, until 50 or more showing a full contour and distinct content were available for analysis in each animal. The varicosities identified as such by their diameter, >0.25 µm, were analyzed by using the public domain IMAGE J processing software from NIH (v.1.45), for the long and short axis and cross-sectional area. They were then categorized as showing or not a synaptic junctional complex, i.e. a localized straightening of apposed plasma membranes associated with a slight widening of the intercellular space and a thickening of the pre-and/or postsynaptic membrane. All synaptic junctions were also characterized as symmetrical or asymmetrical and the length of junctional complexes measured. The synaptic incidence observed in single section was then extrapolated to the whole volume of varicosities by means of the formula of Beaudet and Sotelo ^69^, using the long axis as diameter, according to Umbriaco, Watkins ^70^.

### Statistical analysis

Statistical analyses were performed by using GraphPad Prism 10 (GraphPad Software). Statistical differences between mice with altered VGluT3 expression and control animals were assessed with the non-parametric two-tailed Mann-Whitney U-test.

## Results

### Characterization of the ePet-cre;Cas9-eGFP mouse line

To enable CRISPR-mediated gene editing selectively in 5-HT neurons, we generated a mouse line expressing Cas9 endonuclease and the enhanced green fluorescent protein (eGFP) under ePet-Cre control. To this end, Rosa26-LSL-Cas9 knock-in mice, which have a floxed-STOP cassette upstream of Cas9-eGFP, were crossed with ePet-Cre mice ^62^. For the sake of simplicity, we abbreviate this new mouse line as “ePet-cre;Cas9-eGFP” (Fig. 1A).

**Figure 1.**
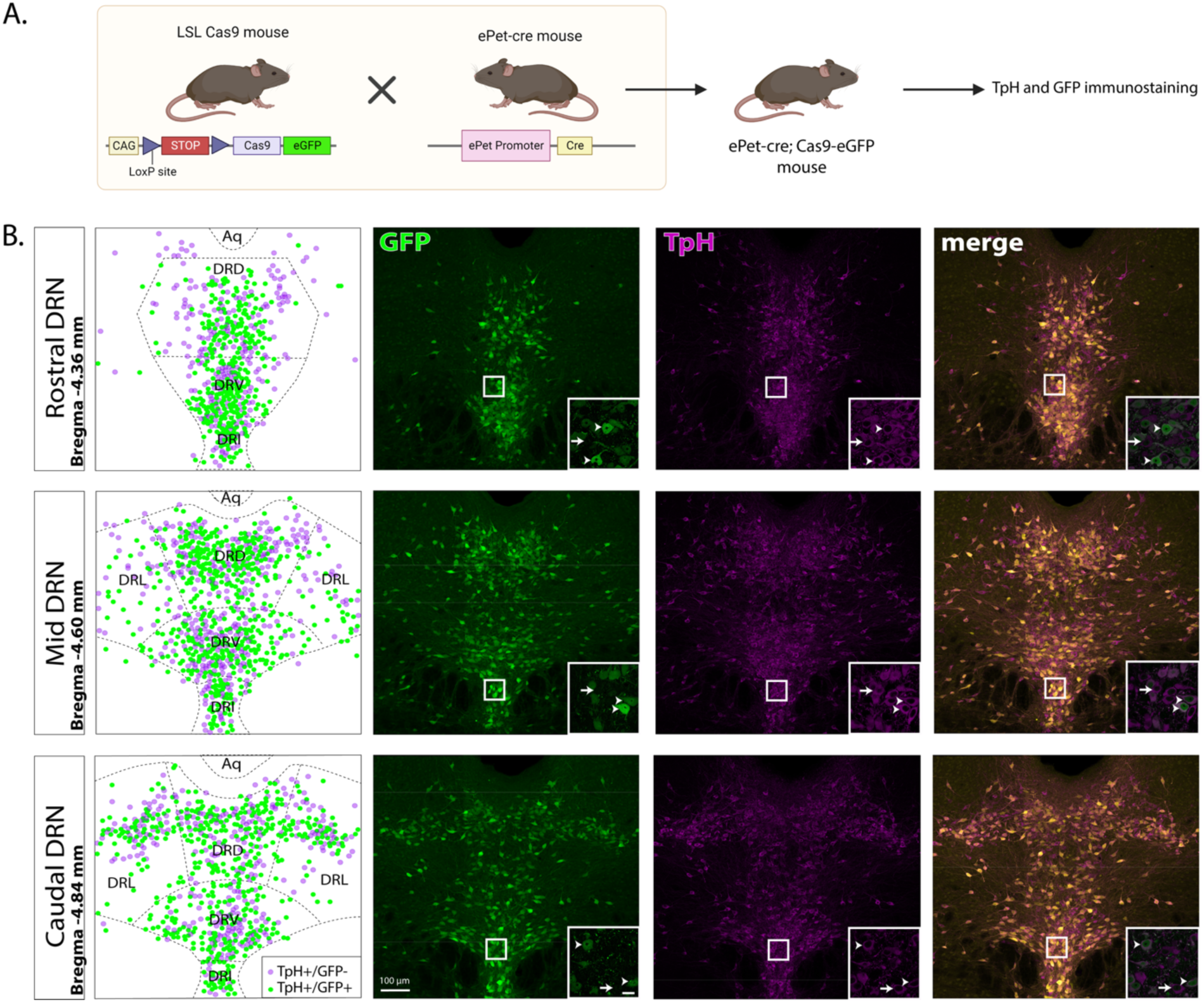
Cas9 and eGFP expression in 5-HT neurons of ePet-cre;Cas9-eGFP mice. (**A**) Schematic of the breeding for the new ePet-cre;Cas9-eGFP mouse line. (**B**) Cas9 and eGFP are expressed in 5-HT neurons of the DRN. TpH (purple) and GFP (green) immunopositive neurons were counted in the rostral (bregma –4.36 mm, upper row), mid (bregma –4.60 mm, mid row) and caudal (bregma –4.84mm, lower row) regions of the DRN. Arrows indicate TpH+/GFP- neurons and arrowheads point to TpH+/GFP+ neurons. Cells immunoreactive for both TpH and GFP (green dots) or only TpH (purple dots) in the DRN are depicted in the schematic illustrations in the left column. DRN, dorsal raphe nucleus; DRD, dorsal part of the DRN; DRV, ventral part of the DRN; DRL, lateral part of the DRN; DRI, interfascicular part of the DRN. Scale bar: 100 µm.

In the resulting ePet-cre;Cas9-eGFP mouse line, all eGFP+ neurons in the DRN co-expressed the tryptophan hydroxylase (TpH) confirming that Cas9 is restricted to 5-HT neurons (Fig. 1B). The quantification indicated that Cas9-eGFP was expressed in the majority of DRN 5-HT neurons, with colocalization rate of 63.9 ± 1.9 % in the rostral DRN, 71.5 ± 4.0 % in the mid-DRN, and 76.7 ± 8.2 % in the caudal DRN, with a smaller fraction of TpH+ neurons lacking eGFP. It should be noted that some TpH+ neurons immunoreactive for GFP were also observed in the MRN (Fig. S2A). Thus, the ePet-cre;Cas9-eGFP lines provides 5-HT-restricted expression of the Cas9 endonuclease and in the majority of DRN 5-HT neurons.

### Efficient and specific knockdown of VGluT3 by CRISPR/Cas9 approach in vitro

We designed two gRNAs against *Slc17a8* (*Vglut3* gene): gRNA1 targeting exon 1 and gRNA2 targeting exon 2 (Table S1). Both gRNAs were expressed from the same plasmid under the U6 promoter. Because the efficacy and specificity of the CRISPR/Cas9 gene editing depends strongly on the sequence of gRNAs ^71, 72^, we first validated their performance *in vitro*. Indels generated from the transfections were analyzed using the TIDE method for CRISPR/Cas9 gene editing ^73^. TIDE analysis revealed minimal activity for gRNA1 (indel frequency of 1.6 %; Fig. S1A), whereas gRNA2 produced a robust editing profile with a 61.9 % indel efficacy (Fig. S1B). The mutations at the gRNA2 site consisted primarily of single-base insertions or deletion. These results indicate that gRNA2 is effective in generating indels in *Slc17a8*. As both gRNAs were packaged into the AAV vector, subsequent *in vivo* experiments employed this dual-gRNA construct, with efficiency editing expected to be driven by gRNA2 to disrupt VGluT3 expression.

### VGluT3 knockdown and overexpression in DRN 5-HT neurons of adult ePet-cre;Cas9-eGFP mice

Using ePet-cre:Cas9-eGFP mice, we applied CRISPR/Cas9 technology to disrupt VGluT3 expression selectively in DRN 5-HT neurons. To do this, adult mice were injected in the DRN with an AAV encoding two gRNAs targeting *Slc17a8* (*Vglut3*), both under the control of U6 promoter, together with a Cre-dependent cassette for mCherry (AAV-U6-gRNA(VGluT3)-DIO-mCherry). To overexpress VGluT3, ePet-Cre mice were injected in the DRN with a Cre-dependent AAV encoding VGluT3 together with mCherry (AAV-DIO-VGluT3-p2A-mCherry). Mice receiving VGluT3-targeting gRNAs are referred to as VGluT3 conditional KO mice and those injected with the overexpression vector, as VGluT3 overexpressing mice. Control animals were injected with an AAV encoding scrambled gRNA (AAV-U6-gRNA(Scramble)-DIO-mCherry). All AAVs expressed mCherry, facilitating detection of infected neurons in the DRN (Fig. 2B). Immunohistochemistry showed expression of mCherry in the vast majority of the TpH+/GFP+ neurons (81.0 ± 7.2 %; Fig. S2A) confirming the effective transduction. In contrast, very few TpH+/GFP+ neurons in the MRN were AAV-infected (mCherry+, 1.3 ± 0.2 %, Fig. S2A), consistent with restricted injection sites.

**Figure 2.**
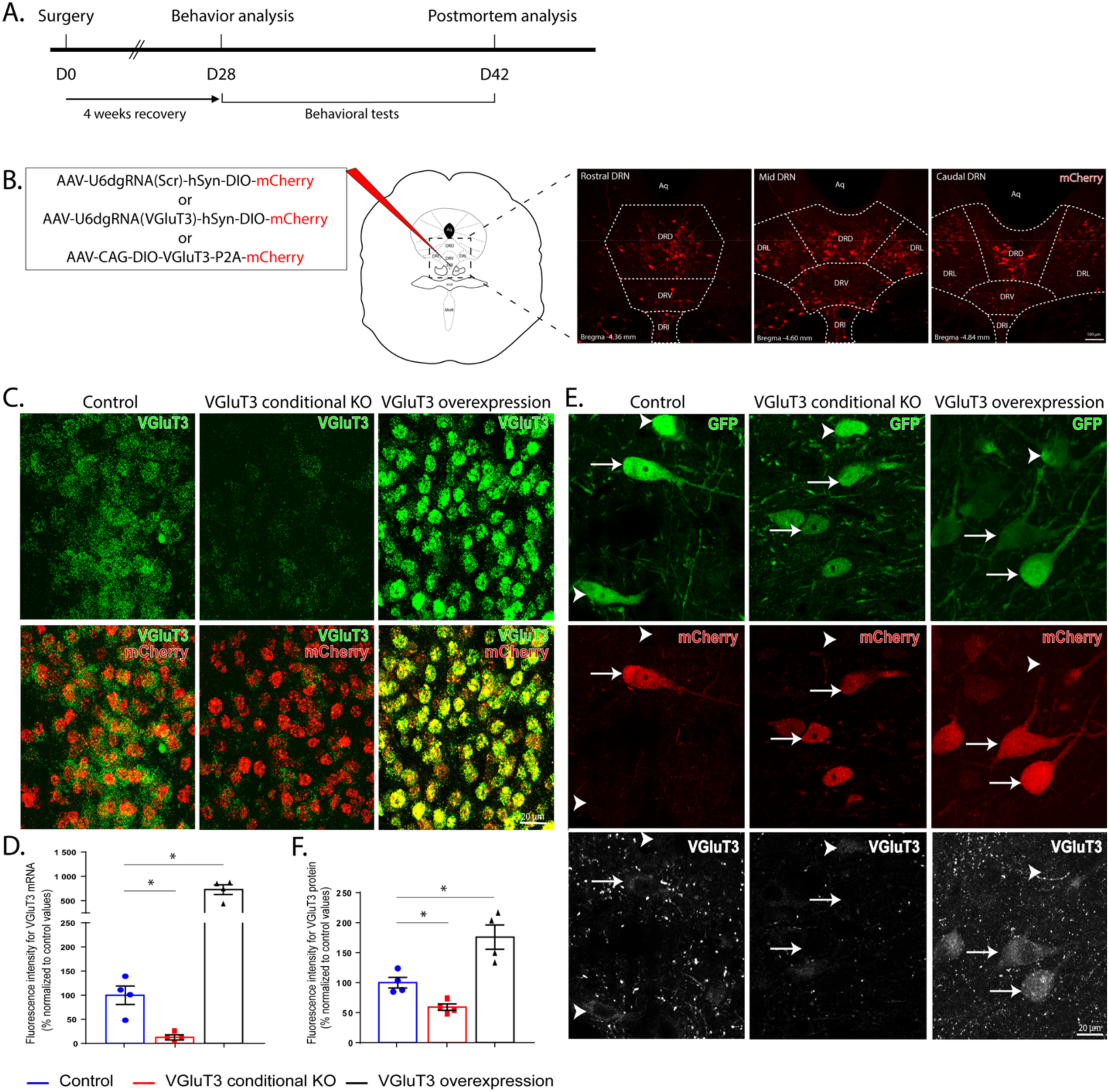
Conditional knockdown or overexpression of VGluT3 is obtained through AAV infection of 5-HT neurons of the DRN. (**A**) Experimental timeline of the viral stereotaxic injection followed by behavioral tests and postmortem analysis. (**B**) Representative viral infection in the DRN. The AAV-infected 5-HT neurons are labeled in red (mCherry+) through the whole rostrocaudal extent of the DRN. The three experimental groups are: control mice injected with AAV-U6-dgRNA(Scramble)-hSyn-DIO-mCherry (AAV-dgScramble), VGluT3 conditional KO mice injected with AAV-U6-dgRNA(VGluT3)-hSyn-DIO-mCherry (AAV-dgVGluT3) and VGluT3 overexpressing mice injected with AAV-CAG-DIO-VGluT3-P2A-mCherry (AAV-VGluT3). (**C**) Confocal microscope image of fluorescent in situ hybridization (RNAscope®) showing VGluT3 (*Slc17a8*, green) and mCherry (mCherry, red) mRNAs in 5-HT neurons of the DRI of AAV-injected mouse. Intense signals for mCherry mRNA were observed in AAV-infected 5-HT neurons of the DRN. Different signal levels for VGluT3 mRNA are observed in these neurons based on the experimental conditions. (**D**) Histogram showing the fluorescence intensity level for VGluT3 mRNA in control, KO and overexpression conditions. Low and high VGluT3 mRNA intensity signals are found in the KO and overexpression conditions, respectively, when compared to control animals. (**E**) Representative immunostaining for GFP (green), mCherry (red) and VGluT3 (white) proteins in the DRI. 5-HT neurons labeled in green were infected either by the AAV-dgScramble (control), by the AAV-dgVGluT3 (VGluT3 conditional KO) or by the AAV-VGluT3 (VGluT3 overexpression), all shown in red (mCherry). Arrows indicate AAV-infected 5-HT neurons and arrowheads AAV-non infected 5-HT neurons. (**F**) Bar graph showing the fluorescence intensity level for VGluT3 protein in control, KO and overexpression conditions. VGluT3 protein intensity signal in DRN 5-HT neurons is lower in KO condition compared to the overexpression condition. Data are represented as means ± SEM. Symbols are individual values obtained for each mouse. Four mice were used in each experimental group. * *P* < 0.05 using the two-tailed Mann-Whitney U test (D, F). Scale bars: 100 μm (B), 20 µm (C, E). DRN, dorsal raphe nucleus; DRD, dorsal part of the DRN; DRV, ventral part of the DRN; DRL, lateral part of the DRN; DRI, interfascicular part of the DRN.

Following AAV injections, transgene expression typically peaks within 2-6 weeks, depending on the AAV capsid ^74–76^. Accordingly, animals were analyzed 4 weeks after injection to ensure stable expression (Fig. 2A). *In situ* hybridization and immunostaining were performed on DRN brain sections to assess editing and overexpression efficiency (Fig. 2C-F). Fluorescent *in situ* hybridization (RNAscope®) detected both VGluT3 and mCherry mRNAs in DRN 5-HT neurons in AAV-injected mice (Fig. 2C). Quantification of VGluT3 mRNA fluorescence intensity within infected 5-HT neuron showed a significant decrease in the KO condition (12.5 ± 5.3 %, *P* = 0.029) and a significant increase in the overexpression condition (728.0 ± 99.8 %, *P* = 0.029) compared with scrambled control condition (99.8 ± 19.0 %, Fig. 2D). In the overexpression condition, we noticed that puncta for VGluT3 mRNAs were mostly located in cell nuclei, in contrast to control condition where they were found in both the cell nuclei and the cytoplasm (Fig. 2C). We next examined protein expression level by immunostaining for eGFP, mCherry and VGluT3 (Fig. 2E). VGluT3 immunoreactivity was significantly reduced in the KO condition (59.3 ± 5.4 %, *P* = 0.029) and elevated in the overexpression condition (176.0 ± 20.3 %, *P* = 0.029) when compared to scrambled controls (100.0 ± 8.9 %, Fig. 2F). Interestingly, a subset of AAV-infected 5-HT neurons in the lateral and dorsal DRN displayed particularly strong VGluT3 labelling in the overexpression condition. Finally, to assess potential toxicity, TpH+ cells counts in the DRN were compared for each experimental group and no significant difference in the number of TpH+ neurons were found between conditions.

### VGluT3 levels do not alter the density of DRN 5-HT axon varicosities in the lateral hypothalamus but influence synaptic profile

After confirming VGluT3 mRNA and protein levels in the DRN, we examined 5-HT axon varicosities in one of their major projection targets, the lateral hypothalamic area (LHA), which exhibits a high density of 5-HT axons arising from the DRN ^50, 77–79^ displaying VGluT3 immunoreactivity ^51^.

Transverse brain sections from control, VGluT3-depleted and VGluT3 overexpressing mice were immunostained for mCherry and VGluT3 to assess whether VGluT3 levels affect the density or morphology of 5-HT axon varicosities (Fig. 3A). The number of mCherry+ terminals and total axon length were quantified in the LHA to obtain the number of axon varicosities per 10 μm of axon. Although not statistically significant, the number of mCherry+ axon varicosities per 10 μm of axon appears slightly higher in VGluT3 overexpressing mice (1.9 ± 0.3 axon varicosity / 10 µm of axon) compared to controls (1.3 ± 0.2, *P* = 0.200, Fig. 3B). VGluT3 immunoreactivity within mCherry+ axon varicosities reflected the expected manipulations. VGluT3 signal was present only in a small fraction of mCherry+ axon varicosities in the KO group (3.8 ± 1.7 %, *P* = 0.029), whereas it was present in nearly all varicosities in the overexpressing mice (97.8 ± 1.3 %, *P* = 0.029), compared to control mice (66.8 ± 9.4 %, Fig. 3C). These findings, consistent with *in situ* hybridization and immunostaining results, confirm efficient VGluT3 depletion and overexpression in respective groups. Morphologically, control 5-HT axons appeared thin and exhibit both fusiform and spherical varicosities. In VGluT3-depleted mice, varicosities were predominantly fusiform, whereas overexpression increased the proportion of round and larger axon varicosities (Fig. 3A).

**Figure 3.**
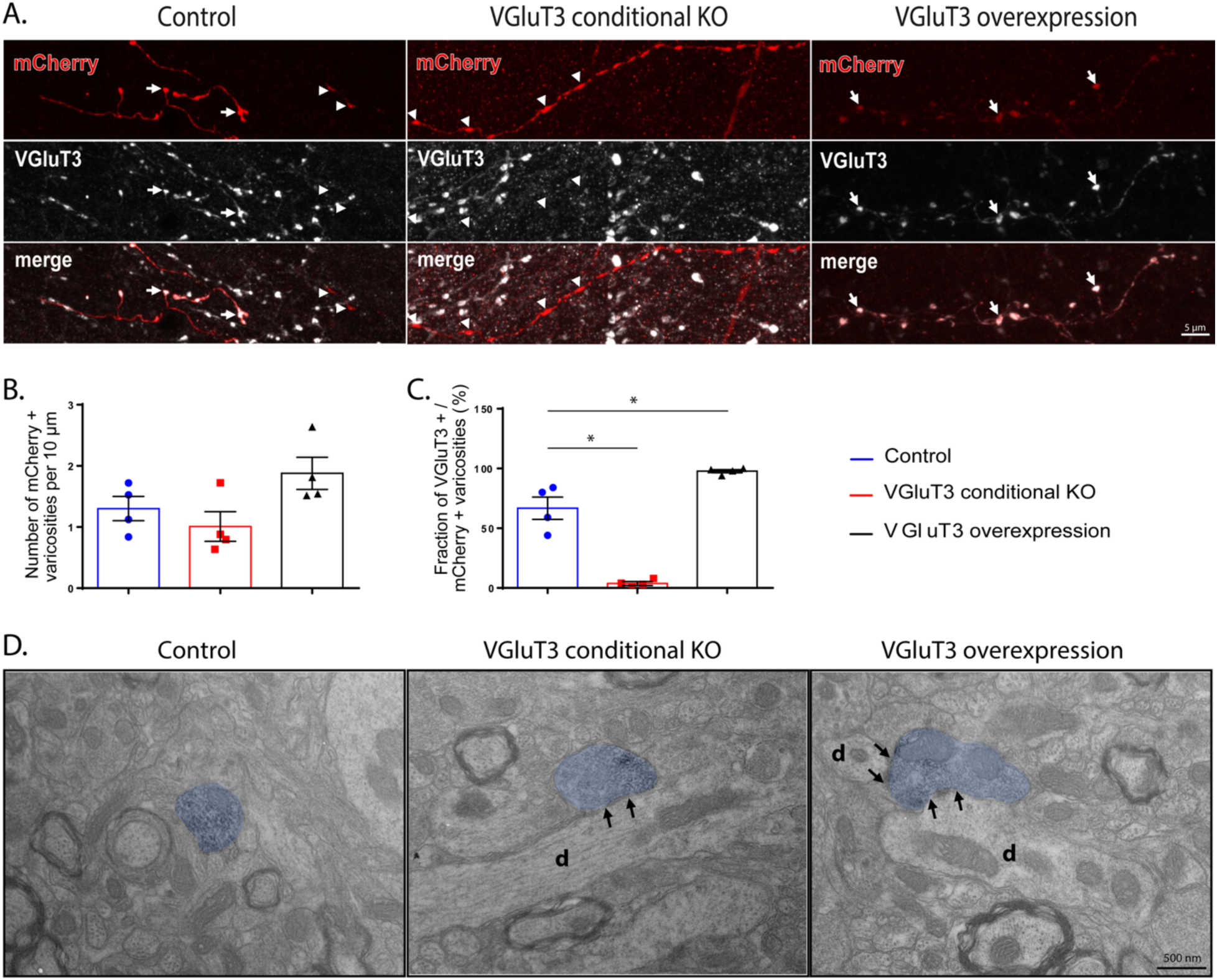
The number and ultrastructural features of LHA 5-HT axon varicosities are not affected by the level of VGluT3. (**A**) Representative immunostaining for mCherry (red) and VGluT3 (white) proteins contained in 5-HT axons found in the LHA of control, VGluT3 conditional KO and VGluT3 overexpressing mice. Arrows indicate mCherry+/VGluT3+ varicosities and arrowheads mCherry+/VGluT3- varicosities. Note the larger size of mCherry+ axon varicosities in the VGluT3 overexpressing mice, compared to VGluT3-depleted mice (**B-C**) Histograms showing the number of mCherry+ varicosities per 10 μm (**B**) and the proportion of mCherry+ varicosities that are VGluT3+ (**C**) in control, KO, and overexpression conditions. The number of 5-HT (mCherry+) varicosities per 10 μm of axon (**B**) is similar between the three experimental groups, although the percentage of mCherry+ varicosities containing VGluT3+ (**C**) is significantly lower in KO condition compared to overexpressing mice. (**D**) Electron micrographs of 5-HT (mCherry+, blue) axon varicosity profiles. The 5-HT axon varicosity profile depicted asymmetric synaptic contacts (between black arrows) with dendritic branches (d) of LHA neurons. Data are represented as means ± SEM. Symbols represent individual values obtained for each mouse. Four mice were used in each experimental group. * *P* < 0.05 using the two-tailed Mann-Whitney U test (C). Scale bars: 5 μm (A) and 500 nm (D).

Based on our confocal observations, we next examined whether changes in VGluT3 expression affected the ultrastructural organization and synaptic relationship of DRN 5-HT axon varicosities in the LHA. As shown on the electron micrographs (Fig. 3D), mCherry+ axon varicosities could be readily identified as their axoplasm was filled with 3, 3’-diaminobenzidine (DAB) immunoprecipitate. Ultrastructural features of these DAB-filled varicosities were quantified across conditions (Table 1). No significant differences were found in most morphological features. However, VGluT3 overexpression was associated with higher proportion of asymmetric synaptic contacts (29.6 ± 1.7 %) compared with controls (20.0 ± 3.5 %, *P* = 0.043). In this group, mCherry+ axon varicosity predominantly formed asymmetric synaptic contacts onto dendritic branches of LHA neurons (Fig. 3D). Altogether, these observations show that VGluT3 manipulation does not substantially alter overall density or size of DRN-derived 5-HT varicosities in the LHA, but influence their synaptic organization, promoting a greater incidence of asymmetric (putative excitatory) contacts under VGluT3 overexpression.

**Table 1.** Ultrastructural features of 5-HT axon varicosities observed in the LHA.

|  | Lateral hypothalamus |  |  |
| --- | --- | --- | --- |
|  | Control | VGluT3 conditional KO | VGluT3 overexpression |
|  | mCherry+ | mCherry+ | mCherry+ |
| Mice | 4 | 4 | 4 |
| Number examined | 291 | 254 | 226 |
| Dimensions |  |  |  |
| Short axis (µm) | 0.62 (±0.04) | 0.70 (±0.03) | 0.69 (±0.03) |
| Long axis (µm) | 1.01 (±0.04) | 1.17 (±0.07) | 1.05 (±0.05) |
| Aspect ratio | 0.64 (±0.03) | 0.63 (±0.02) | 0.68 (±0.02) |
| Diameter (µm) | 0.82 (±0.04) | 0.94 (±0.05) | 0.87 (±0.04) |
| Area (µm<sup>2</sup>) | 0.54 (±0.07) | 0.69 (±0.06) | 0.62 (±0.06) |
| % with mitochondria | 35.1 (±5.1) | 32.9 (±5.7) | 36.1 (±3.9) |
| Synapse |  |  |  |
| Asymmetric (%) | 20.0 (±3.5) | 41.0 (±9.3) | 29.6 (±1.7)* |
| Symmetric (%) | 80.0 (±3.5) | 59.0 (±9.3) | 70.4 (±1.7)* |
| Synapse length (µm) | 0.36 (±0.03) | 0.41 (±0.05) | 0.42 (±0.03) |
| Extrapolated synaptic incidence (%) | 28.8 (±9.4) | 48.4 (±9.1) | 38.5 (±4.0) |

Mean ± SEM from four mice
\* $P < 0.05$ for VGluT3 overexpression versus control
Statistical analysis was performed using Mann Whitney test.

### VGluT3 depletion in DRN 5-HT neurons induces anxiety-like behaviors in adult mice

Four weeks following AAV injections, the three experimental groups were tested for motor and non-motor behaviors to assess the impact of VGluT3 depletion or overexpression on locomotion, anxiety and depressive-like behaviors (Fig. 4A).

**Figure 4.**
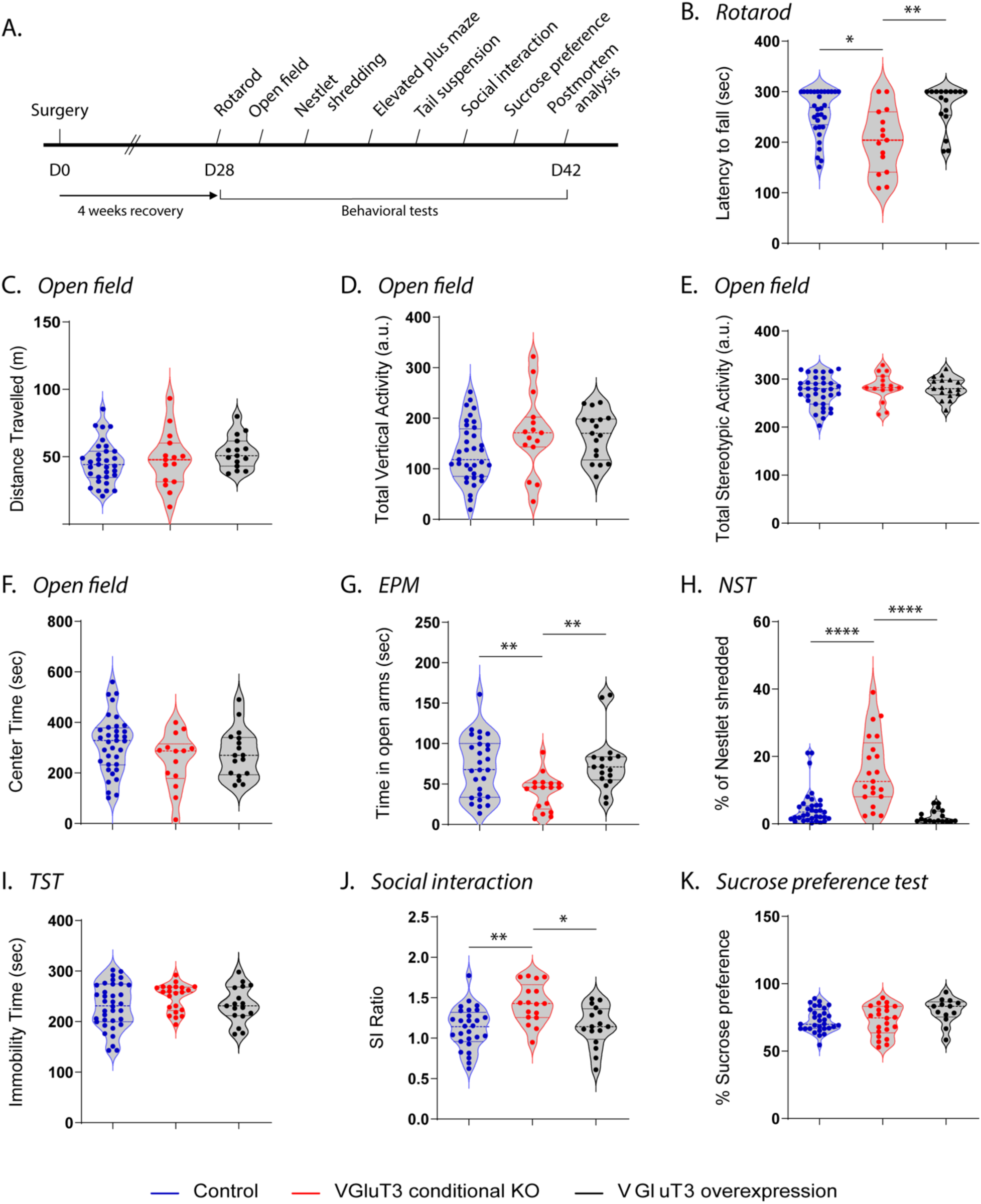
Behavioral assessment of VGluT3*-*depleted and overexpressing mice. (**A**) Experimental timeline showing viral injection (surgery) followed by behavioral tests used to assess the effect of VGluT3 depletion or overexpression on motricity, anxiety and depressive-like behaviors of ePet-cre;Cas9-eGFP mice. (**B-K**) Violin plots of data distributions (probability density), median and interquartile ranges for the different behavioral tests. Coordination, endurance, and muscular strength as assessed with the rotarod (**B**) were affected in VGluT3 conditional KO mice which fell faster than control (*P* = 0.004). No difference in the spontaneous motor activity (distance travelled, vertical and stereotypic activity; **C-E**) as assessed in the open field was noted between the three experimental groups. Anxiety-like behaviors were scored in the open field (**F**), nestlet shredding test (NST, **G**) and elevated plus maze (EPM, **H**). VGluT3 conditional KO mice tend to spend less time in the center of the open field (*P* = 0.1) and significantly less time in the open arms of the EPM compared to control mice (*P* = 0.005). Nest building activity was significantly increased in VGluT3 conditional KO mice (*P* < 0.0001), whereas it was significantly reduced in VGluT3 overexpressing mice (*P* = 0.038) when compared to control animals. Assessment of depressive-like behaviors was done using the tail suspension test (TST, **I**), social interaction test (**J**) and sucrose preference test (**K**). No depressive-like behavior was observed in either VGluT3-depleted or VGluT3 overexpressing mice. However, VGluT3 conditional KO mice displayed increased social interactions (*P* < 0.0005), while VGluT3 overexpressing mice showed a greater preference for sweetened water (*P* = 0.023). Dots represent individual values obtained for each mouse. * *P* < 0.05; ** *P* < 0.005; *** *P* < 0.001; **** *P* < 0.0001 using the two-tailed Mann-Whitney U test.

Motor coordination and endurance were assessed using the rotarod test (Fig. 4B). VGluT3 conditional KO mice fell significantly faster (203.4 ± 16.2 s) than controls (257.7 ± 8.3 s, *P* = 0.004). Spontaneous motor activity including distance travelled, vertical and stereotypic activity was assessed in the open field (Fig. 4C-E) and no significant differences were noted between the three experimental groups. Anxiety-like behaviors were examined in the open field (Fig. 4F), the elevated plus maze (EPM, Fig. 4G) and the nestlet shredding test (Fig. 4H). Avoidance of the center in the open field or of open arms in the EPM reflects anxiety-like behavior, whereas increased nestlet shredding indicates stress-induced repetitive compulsive behavior. VGluT3 conditional KO mice tended to spend less time in the center of the open field (249.4 ± 29.0 s) than controls (312.6 ± 19.4 s; *P* = 0.114), and showed increased vertical activity in the periphery (170.8 ± 20.2 vs. 131.6 ± 9.9; *P* = 0.094), suggesting a trend toward increased anxiety. In the EPM, VGluT3 conditional KO mice spent significantly less time in the open arms (40.1 ± 5.3 s) than control mice (71.5 ± 7.2 s, *P* = 0.005). Moreover, nestlet-shredding activity was significatively increased for VGluT3 conditional KO mice (15.6 ± 2.3 %, *P* < 0.0001) and significatively decreased for VGluT3 overexpressing mice (2.3 ± 0.5 %, *P* = 0.043), compared to controls (4.8 ± 0.9 %). Depressive-like behavior was assessed using the tail suspension (Fig. 4I), social interaction (Fig. 4J) and sucrose preference (Fig. 4K) tests. No significant depressive-like phenotype was detected in VGluT3 conditional KO or VGluT3 overexpressing mice. However, VGluT3 conditional KO mice exhibited significantly greater social interaction with the target mouse (1.4 ± 0.1), compared to controls (1.1 ± 0.1, *P* = 0.0005). Moreover, VGluT3-overexpressing mice showed increased in sucrose preference (80.0 ± 2.6 %) when compared to control animals (73% ± 1.5%, *P* = 0.023).

Collectively, these results indicate that manipulation of VGluT3 expression in DRN 5-HT neurons influence anxiety-related behaviors, with depletion enhancing anxiety-like and repetitive behaviors, and overexpression producing opposite or mild anxiolytic effects. These findings suggest that VGluT3 contributes to the modulation of emotional behaviors by DRN 5-HT neurons.

## Discussion

In the present study, we assessed VGluT3 functions by either depleting VGluT3 using the CRISPR/Cas9 method, or overexpressing this gene, specifically in DRN 5-HT neurons of adult mice. We hypothesized that altered VGluT3 expression by these neurons could impact synaptic connectivity and consequently, behaviors. In line with this hypothesis, we showed that (1) overexpression of VGluT3 by DRN 5-HT neurons increased the proportion of asymmetric synapses formed by their terminals on LHA neurons and that (2) depletion of VGluT3 in DRN 5-HT neurons was associated with anxiety-like behaviors and alterations in motor performance and social interactions. Below, we discuss these findings in the light of relevant literature.

The 5-HT neurons spatially distributed along the rostro-caudal axis of the brainstem within raphe nuclei originally designated as groups B1–B9 by Dahlstrom and Fuxe ^80^. The DRN (groups B6 and B7) contains the vast majority of 5-HT neurons in the central nervous system ^44^ and provides most of the ascending 5-HT innervation of the midbrain and forebrain ^81^. In mice, approximately two-thirds of the DRN 5-HT neurons express VGluT3 ^48^, and this proportion is even higher for 5-HT neurons located along the midline ^48, 51, 82^. VGluT3 is also expressed by most of the 5-HT neurons of the MRN ^8, 9, 13–17^, as well as by other type of neurons located in the cerebral cortex ^8, 9^, the hippocampus ^10, 11^, the striatum ^9, 12–14^, and the peripheral nervous system ^18–21^. The broad distribution of VGluT3 expressing neurons complicates efforts to characterize VGluT3 functions exclusively to DRN 5-HT neurons. Moreover, constitutive knockout or overexpression models induce lifelong alterations in gene expression that may trigger compensatory mechanisms, precluding a precise assessment of VGluT3 functions in 5-HT neurons.

To study VGluT3 function in a specific population of 5-HT neurons, we generated a transgenic mouse line, ePet-cre:Cas9-eGFP, allowing spatially and temporally controlled gene editing in 5-HT neurons of adult mice. In this transgenic mouse line, approximately 70 % of TpH+ neurons in the DRN expressed eGFP+, reflecting Cre-dependent activation of the Rosa-LSL-Cas9-eGFP allele in *Pet-1*-positive 5-HT neurons (Fig. 1B). Previous characterizations of the ePet-cre mice demonstrated that Cre expression is largely restricted to 5-HT neurons ^62, 83^, although the exact proportion of labeled cells was not quantified. Our observation is therefore consistent with prior evidence that *Pet1* is expressed in most, but not all, 5-HT neurons ^84, 85^. This partial labeling likely reflects the known developmental heterogeneity of 5-HT populations. Indeed, in *Pet1*- ⁄-mice, approximately 70 % of raphe neurons failed to differentiate into 5-HT neurons ^84, 85^, and conditioned deletion of *Pet1* in adult mice resulted in > 70 % reduction in Pet1 mRNA expression in the groups B5–B9 of the raphe nuclei, and a 50 % decrease of *TpH* expression in the rostral clusters ^86^. These findings indicate that, while *Pet1* is a critical transcription factor required for the specification of most 5-HT neurons, a subset can maintain 5-HT phenotype independently of *Pet1*, possibly through compensatory developmental mechanisms involving Shh, Fgf4/8, Nkx2.2, Nkx6.1, Gata2, Lmx1b, Hmx, En and Pet1 ^83, 87–89^. Our quantitative analysis also indicated that more than 80 % of TpH+ DRN neurons were infected by our AAVs, indicating high but incomplete viral coverage. Together, these results confirm that VGluT3 manipulation in our models was largely restricted to DRN 5-HT neurons, while incomplete Cre expression likely limited the magnitude of the observed effects.

We confirmed that our VGluT3 KO and overexpression strategies resulted in significant reduction and increase in VGluT3 mRNA and protein levels, respectively. Interestingly, while VGluT3 mRNA levels increased approximately 10-fold in the overexpression condition compared to the control, the corresponding VGluT3 protein levels showed only a ∼2-fold increase (Fig. 2D, F). We also observed that VGluT3 mRNAs puncta detected by RNAscope were frequently concentrated in the cell nuclei. Altogether, these observations suggest an effective post-transcriptional regulatory mechanism for VGluT3 production (for review, see ^90^), potentially through various factors including microRNAs like miR-137 ^91^. A tight regulation of VGluT3 level might be particularly important to limit excitotoxicity. While overexpression of VGluT transporters in “pure” glutamatergic neurons can induce excitotoxicity ^92^, we did not observe any 5-HT cell loss in the DRN following overexpression and no evidence of excitotoxicity in downstream target regions, including the LHA, suggesting that moderate VGluT3 overexpression in 5-HT neurons is well tolerated.

The potential ability of DRN 5-HT neurons to release both 5-HT and glutamate may have important implications for synaptic plasticity and neuroadaptive mechanisms. This may be particularly important for DRN neurons considering the possibility of a complex trafficking mechanism of different synaptic vesicles along their highly collateralized axon ^49^. This would offer the ability for 5-HT neurons of the DRN to co-release glutamate along with 5-HT in different proportion according to their multiple target sites. Previous electron microscope studies have revealed that a large proportion of 5-HT axon varicosities are devoid of any synaptic contact and that such low synaptic incidence may vary depending on target sites ^93^. In contrast, “pure” glutamatergic systems labeled with VGluT1 or VGluT2 have often been reported to be entirely synaptic, each axon varicosity being endowed with more than one synapse ^94, 95^. Because the proportion by which 5-HT axon varicosities contain VGluT3 has also been shown to vary according to target sites ^49^, we wanted to assess if higher VGluT3 content in 5-HT axon terminals could favor the establishment of synapses or contribute to the maintenance of synaptic integrity.

According to original descriptions of 5-HT axons ^96^, our previous study reported that DRN neurons are endowed with two types of axons: (1) axons with fusiform axon varicosities and (2) axons bearing large and spherical axon terminals. We also showed that 5-HT axon varicosities containing VGluT3 are larger than those devoid of this vesicular transporter ^49^. Our confocal microscope observations indicate spherical and larger 5-HT axon varicosities in the LHA of VGluT3 overexpressing mice, compared to fusiform axon terminals observed in VGluT3-depleted mice (Fig. 3A). At the electron microscope level, such differences in shape and size did not reach statistical significance, likely due to a limited number of axon varicosity profiles examined, and to the fact that they were observed from single ultrathin and non-serial sections (Table 1). It should however be noted that a higher aspect ratio was obtained for VGluT3 overexpressing mice, indicating spherical axon varicosities. Since it has previously been reported that larger 5-HT axon varicosities establish more synapses ^97^, we hypothesized that 5-HT axon varicosities from VGluT3 overexpressing mice would show more synaptic contacts than terminals that are devoid of this vesicular transporter. In the LHA of control mice, we found the proportion of 5-HT axon varicosities endowed with a synaptic junction to be low (28.8 %) suggesting the existence of a diffuse mode of transmission for 5-HT in this brain region, in addition to a synaptic mode. Such low synaptic incidence for 5-HT axon varicosities found in the LHA is in agreement with previous reports ^33, 98^. In contrast to our initial hypothesis, depletion or overexpression of VGluT3 by 5-HT neurons of the DRN did not significantly affect the synaptic incidence of 5-HT axon varicosities found in the LHA. In our knockdown model, incomplete loss of VGluT3 suggests that residual protein may be sufficient to maintain synaptic organization. Notably, the number of synaptic vesicles as well as their docking at the plasma membrane was found to be unaffected by a complete loss of VGluT3 in *Vglut3*-/- mice ^21^. Our electron microscope observations of the LHA indicate a significant increase in the proportion of asymmetrical synapses on 5-HT axon varicosities of VGluT3 overexpressing mice. These data suggest that increased VGluT3 levels may bias synaptic morphology toward more asymmetric synapses (excitatory-like), in agreement with previous studies that have shown that 5-HT axon varicosities containing VGluT3 establish mostly asymmetrical synapses on VTA dopamine neurons ^56^ and on sympathetic preganglionic neurons^99^.

At the behavioral level, a lack of consensus remains regarding the primary functions of the DRN 5-HT system, despite an extensive body of literature. Studies on DRN 5-HT neuron functions have reported roles in aversive processing ^100^, reinforcement and reward processing ^54, 55, 101–103^, anxiety-like behaviors ^104^ and locomotion ^105, 106^. Such functional diversity may stem from the fact that DRN 5-HT neurons receive a broad range of afferent projections ^51^ and possess highly collateralized axons that may support projection-dependent glutamatergic signaling, across multiple target sites ^49^.

The specific role of DRN 5-HT neurons in motor behavior appears controversial for review, see ^107^. Previous studies using optogenetics have shown that acute activation of DRN 5-HT neurons induces inhibition of locomotion ^54^ and that such inhibition is context-dependent and does not involve any loss in motor coordination ^106^. In the present study, no significant effect of VGluT3 depletion or overexpression was noted on spontaneous locomotion activity, indicating that the role of 5-HT released from DRN neurons on locomotion may prevail over glutamate co-released by these neurons. It should however be kept in mind that VGluT3 positively modulates 5-HT transmission ^27, 33^ through a mechanism called vesicular synergy, in which glutamate uptake in synaptic vesicles promotes 5-HT storage by increasing the pH gradient that drives vesicular monoamine transporter 2 (VMAT2) ^26^. Therefore, the behavioral consequences and morphological changes observed in our animal models might be directly induced by altered glutamate release by 5-HT DRN neurons but might also partially represent a consequence of changes in 5-HT transmission.

In the present study, motor performance as assessed on the rotarod was significantly decreased in VGluT3-depleted mice. These results are at odds with a previous study conducted in *Vglut3*-/- mice that has shown normal righting reflex and performance on the rotarod ^21^. Such discrepancy may be explained by compensatory mechanisms occurring in *Vglut3*-/- mice or by the cell-type specificity and adult onset of VGluT3 depletion in our model.

The 5-HT system is known to be involved in anxiety ^84^, and its dysfunction has often been associated with mood and anxiety disorders ^108^. Over the years, animal models have been used to demonstrate the role of 5-HT in anxiety. For instance, 5-HT-depleted mice and conditional *Pet-1* KO mice have shown higher anxiety level ^86, 109^. Increased anxiety and depressive-like behaviors have also been correlated with reduced DRN 5-HT neuronal activity ^104^. Consistent with these findings, *Vglut3*-/- mice display anxiety-related phenotype that might partly result from lower 5-HT transmission ^27^. Building on this, our selective VGluT3 depletion in DRN 5-HT neurons increased anxiety-like and repetitive behaviors (EPM, nestlet shredding), whereas overexpression produced opposite effects on shredding. These results indicate that VGluT3 level modulate anxiety-related behavior, possibly through altered 5-HT vesicular function and transmission. It would be of interest to examine 5-HT axon morphology and synaptic incidence in regions, other than the LHA, involved in anxiety, such as the basolateral amygdala ^110^.

Previous studies of depressive-like phenotypes conducted in animal models with altered 5-HT transmission are more controversial. For instance, *Tph2* KO mice showed both antidepressive-like and prodepressive-like behaviors, depending on the behavioral test used ^111, 112^. The *Pet1*-/-mice showed no depressive-like phenotype, as assessed with the tail suspension and forced swim tests ^113^. Likewise, the *Vglut3*-/- mice did not show any despair-like behavior when submitted to the tail suspension test ^27^. In agreement to these reports, our data indicate no change induced by VGluT3 depletion or overexpression on depressive-like behaviors. Stress exposure being an important risk factor for the development of depressive-like disorders, it would be of interest to expose our animal models to chronic stress, before the assessment of depressive-like behaviors.

5-HT transmission is known to be involved in aggressive behavior, in both human and rodents for review, see ^114^. Studies conducted using genetic mouse models of 5-HT depletion such as *Tph2*-/- ^115^ or *Pet1*-/- ^84^ mice reported increased aggressive behavior. Study conducted with *Vglut3*-/- mice did not show any aggressive behavior or social avoidance ^27^. In contrast, VGluT3-depleted mice in our study showed greater social interaction time, which we interpret as increased social approach rather than aggression. Because the nestlet shredding test does not directly evaluate aggression, further behavioral assays, such as resident-intruder, would be required to assess aggression.

In addition to the VTA, nAcc, ventral pallidum, anterior cingulate cortex and orbitofrontal cortex, the DRN has emerged as an important hub in reward circuits, with its 5-HT neurons playing a significant role ^116–119^. Optogenetic studies in mice have shown that acute activation of DRN 5-HT neurons, particularly those projecting to the VTA, produces strong reinforcement signals and enhances reward waiting ^55, 56, 120^. Other studies rather argued that the rewarding effects of the DRN are attributed solely to VGluT3 glutamate signaling ^121, 122^. In the present study, we did not observe any significant differences between VGluT3-depleted mice and control animals during the sucrose preference test, but VGluT3 overexpressing mice showed increased sucrose preference. This finding suggests a potential enhancement of hedonic behavior, though direct modulation of reward circuitry would require further testing using operant or conditioned placed preference paradigms.

The co-transmission of glutamate by monoamine neurons is emerging as an important concept in understanding their functions (for review, see ^123^). Our findings indicate that VGluT3 levels in DRN 5-HT neurons influence synaptic morphology and are associated with distinct behavioral effects, including increased anxiety-like and repetitive behaviors following VGluT3 depletion, and reduced repetitive behavior following overexpression. These results support a modulatory role of VGluT3 in 5-HT circuit structure and function, with behavioral consequences that may arise from altered 5-HT and glutamatergic signaling balance.

## Acknowledgements

The authors would like to thank the CERVO Centre Molecular Tool Platform (https://neurophotonics.ca/fr/pom) for the production of the viral vectors used. This study was supported by research grants from the Canadian Institutes of Health Research (CIHR PJT 470155 to M.P. and 451548 to M.L.) and the Natural Sciences and Engineering Research Council of Canada (CG095382) to M.P., by Parkinson Canada (to M.P. and C.D.P.) and the Canada Foundation for Innovation (CIF 42160 to M.P). M.P., C.D.P and M.L. benefited from a career award from the Fonds de Recherche du Québec-Santé (FRQ-S, 268922; 34974).

## Author contributions

L.S., C.D.P. and M.P. designed the experiments of the study. V.R., S.S.R., E.M. and M.L. contributed to the conception and validation of gRNAs. L.S. performed all surgeries, fluorescent imaging, in situ hybridization and immunohistochemistry experiments, behavioral experiments and data analysis. L.S. performed also electron microscopy experiments, acquisition and analysis (with M.J.W.’s help). L.S. prepared the manuscript with edits by C.D.P., M.L. and M.P. All authors approved the final version.

## Data Availability

The authors confirm that the data supporting the findings of this study are available within the article. Raw data are available from the corresponding author on request.

---

**Supplementary figure 1.**
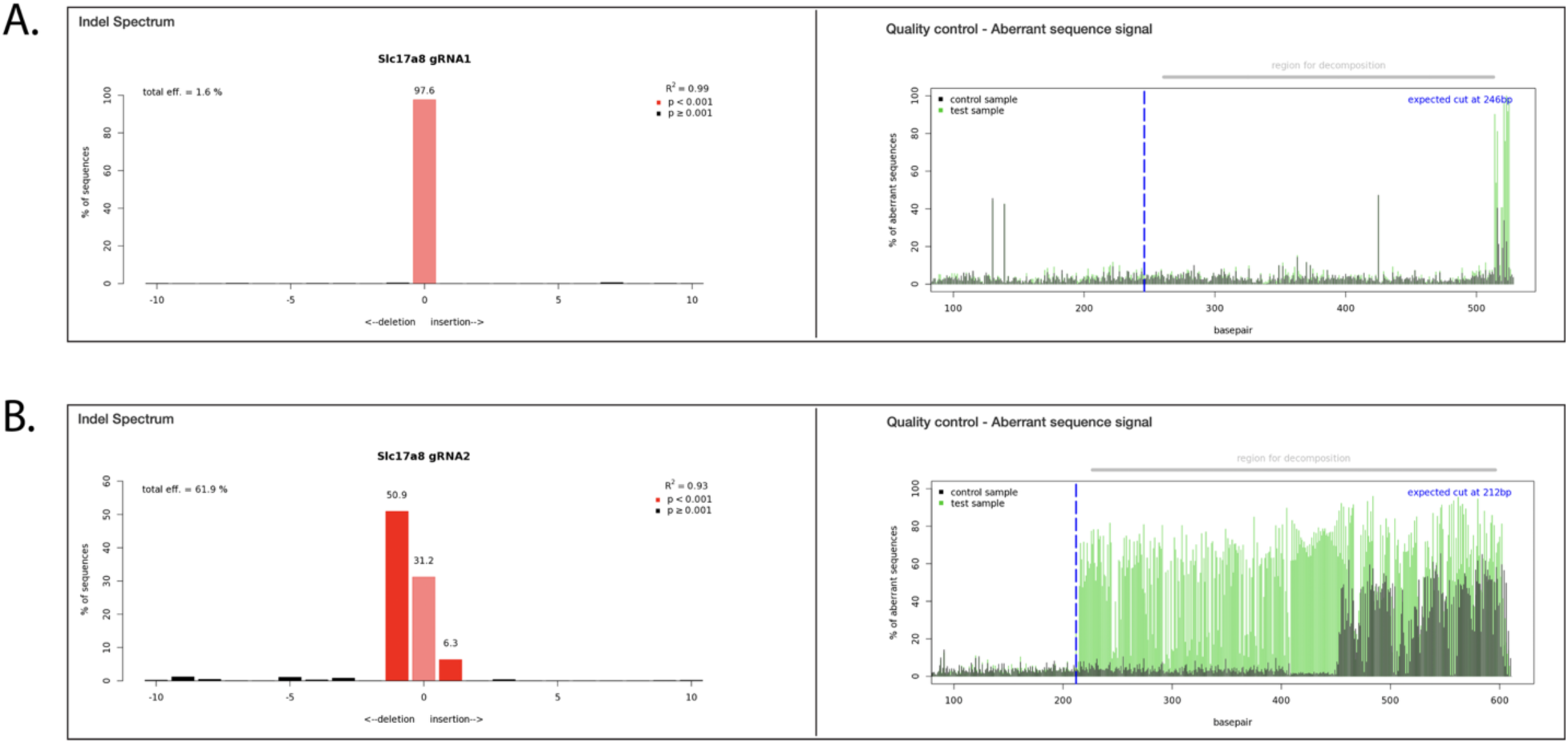
TIDE analysis. (**A-B**) Indels generated from the co-transfection of NIH-3T3 cells with the plasmids encoding for gRNAs used in the present study were analyzed using TIDE method for CRISPR/Cas9 gene editing. Indel spectrum and quality control for gRNA1 and gRNA2 are shown in **A** and **B**, respectively.

**Supplementary figure 2.**
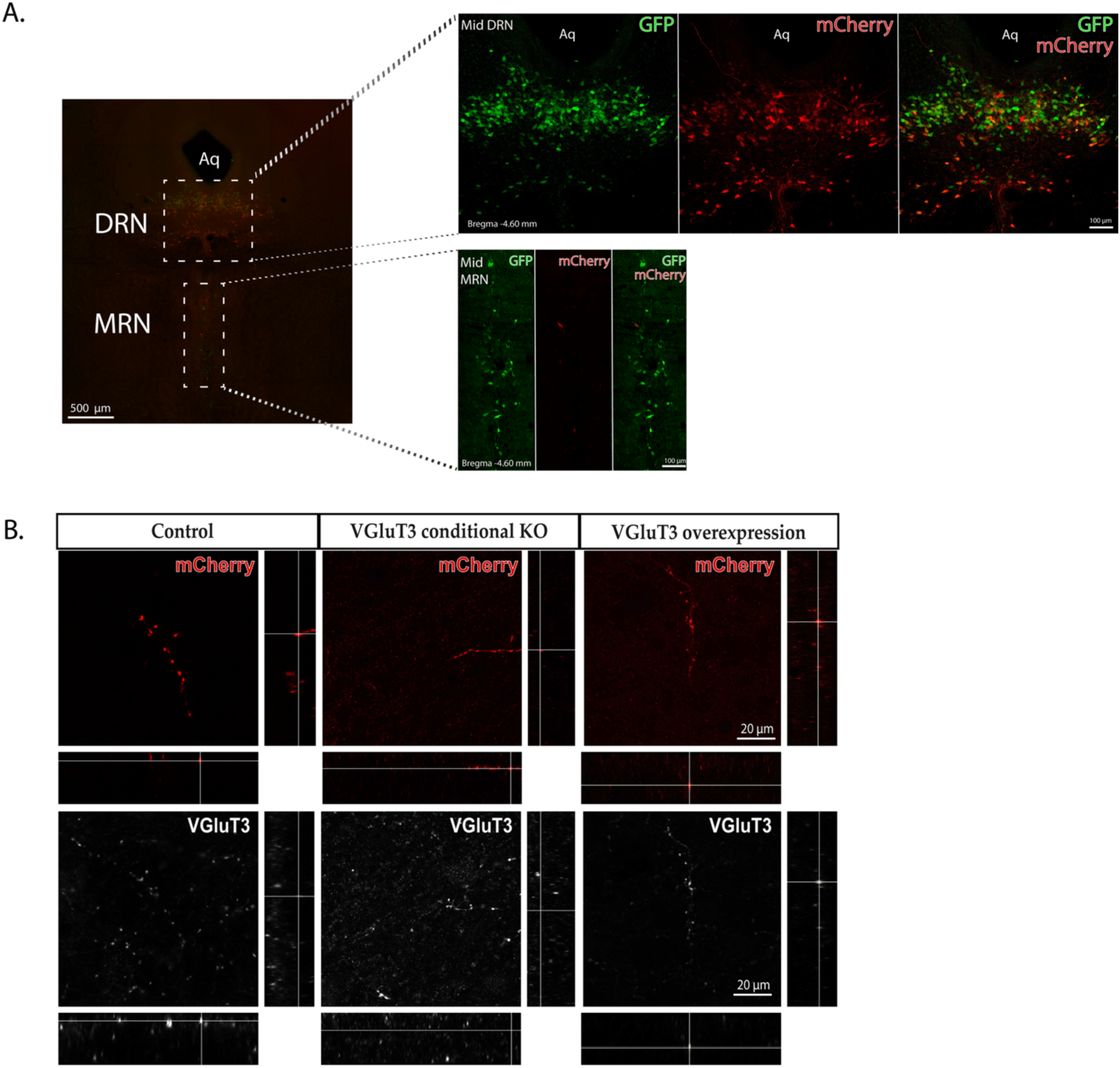
(**A**) Distribution of 5-HT infected neurons expressing mCherry (red) in the DRN and the MRN following intracerebral injection of AAV placed in the DRN of ePet-cre;Cas9-eGFP mice. (**B**) Orthogonal views of axons (mCherry+, red) arising from 5-HT infected neurons of the DRN showing VGluT3+ (white) axon varicosities in the lateral hypothalamic area of control, VGluT3 conditional KO and VGluT3 overexpressing mice.

**Supplementary table 1.**
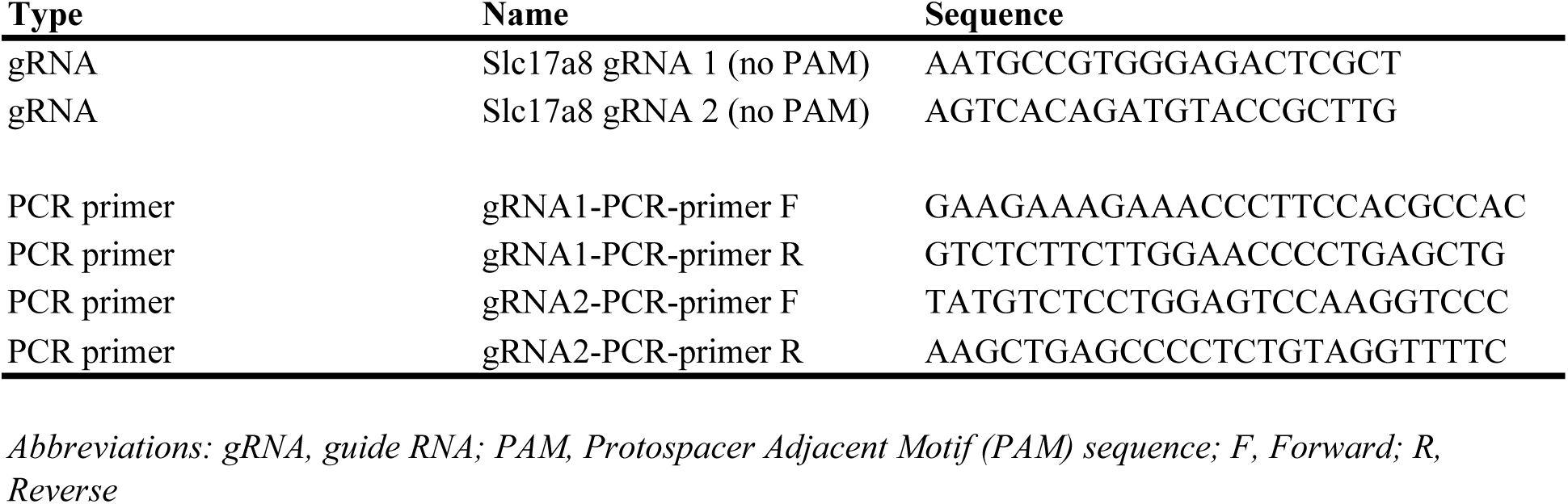
Oligo sequences used (5’-3’)

